# Private forest reserves in Brazil’s agricultural landscapes effectively sustain the occupancy and abundance of multiple threatened large vertebrates

**DOI:** 10.64898/2026.10.16.757138

**Authors:** Joshua P. Twining, Ennio Painkow Neto, Gonzalo Barquero, Kenneth Kellner, José Manuel Vieira Fragoso

**Author notes:** Corresponding author address: Department of Fisheries, Wildlife, and Conservation Science, Oregon State University, 126 Nash Hall, Corvallis OR, 97331.

## Abstract

1. Understanding the factors that shape wildlife abundance and distributions is at the core of applied ecology and wildlife management. Accurate predictions on the population-level outcomes of land management are especially urgent in biodiversity hotspots around the world that are undergoing rapid transformation. In South America, both the Cerrado and Amazon are increasingly being converted to agricultural land-uses to produce soybean and corn and raise cattle. Over the last several decades Brazil has become one of the leading global producers of these products. At the same time, Brazil has some of the strongest federal regulations focused on protecting native habitat. The Native Vegetation Protection Law (or the ‘Forest Code’) requires private properties retain native vegetation as Legal Reserves, and as a result more than half of the country’s remaining native habitat now lies on private lands. Whether retained habitat under this ambitious policy sustains populations of large, wide-ranging vertebrates across entire agricultural production landscapes remains untested.
2. Here, we conducted a landscape-scale survey using camera-trap stations across ∼606,000 km² of Mato Grosso, Brazil’s largest producer of cattle, soybean and corn. We sampled 115 independent sites between November 2024 and September 2025, spanning both crop seasons. We estimated occupancy of jaguars and expected abundance of white-lipped peccary and lowland tapir, using a combination of occupancy and Royle–Nichols models. We fit an abundance-mediated interaction model to formally test the role of prey abundance in driving jaguar occupancy across the landscape.
3. We found high occurrence and expected abundance of all three species. Forest cover was the only covariate common to all top models for every species: predicted jaguar occupancy probability rose as forest cover increased and expected peccary and tapir abundance also increased as a function of forest cover.
4. Our empirical estimates from abundance-mediated interaction models were consistent with jaguar occupancy being a function of both habitat availability and prey (peccary) abundance.
5. These correlational, single-year results demonstrate that native forest retained on private land in one of the world’s most intensively farmed regions are associated with the persistence and local abundance of three threatened large mammals on a landscape-scale.

## Introduction

Land use change is the dominant driver of biodiversity loss and the increasing extinction rates across taxa that define our current epoch (Cardinale et al. 2012; Maxwell et al. 2016; Jaureguiberry et al. 2022). To feed growing human populations and maintain socioeconomic development, humans are continuously expanding the amount of land dedicated to agricultural production (Macdonald *et al*. 2018), along with increasing intensification of the management of existing agricultural land (Balvanera et al. 2019). This pressure on biodiversity resultant from expansion of land-used for cropping and animal husbandry is most acute within developing nations that are also global biodiversity hotspots, e.g., Sundaland and Indo-Burma in southeast Asia, and the Cerrado in South America (Kong et al. 2021). Nonetheless, the challenge of minimizing biodiversity loss while keeping up with production demands for humans is faced globally.

In response to mounting evidence on the impacts of conversion of native habitat to production landscapes increasing biodiversity loss, many countries have put in place policies and regulations to attempt to reduce land conversion to maintain habitat for wildlife (Kremen & Merenlender, 2018). Examples range from species-focused protections such as the Endangered Species Act in the United States and the Species at Risk Act in Canada (Taylor et al. 2005; Hebblewhite, 2017), through habitat-focused safeguards such as plantation regulations and certification schemes of Southeast Asia (Luke et al. 2019), to area-based set-asides such as the establishment of Ecological Focus Areas in Europe (Pe’er et al. 2017) and the Ecological Redline Policy in China (Bai et al. 2018). One of the most ambitious habitat-safeguarding policies in the world is Brazil’s Native Vegetation Protection Law (Law 12,651/2012, also known as ‘the Forest Code’). This law requires every rural property to maintain a Legal Reserve of native vegetation, with the legal reserve requirement varying by habitat and region. Within the Legal Amazon, an administrative region encompassing nine states, 80% of native vegetation must be maintained in forest, and 35% in Cerrado savanna. Outside of the Legal Amazon, this drops to 20% of native habitat. As more than half of Brazil’s remaining native vegetation is on private land, these provisions are arguably as important for maintaining biodiversity in the region as the protected area network (Metzger et al. 2019). Existing empirical research suggests that riparian buffers and private reserves retain much of the medium- and large-bodied mammal and bird assemblage seen in protected areas (Lees & Peres, 2008; Negrões et al. 2011), however, nearly all available evidence stems from studies of individual fragments or corridors on the scale of 10s or 100s of km^2^. The success of the Forest Code policy and the habitat retained under these regulations in sustaining populations of large, low-density, and wide-ranging vertebrates across entire production landscapes (the scale at which policy operates, and species populations persist or decline) remains untested and thus uncertain.

Testing the outcomes of national conservation policy requires estimating wildlife population parameters pertaining to occurrence and population size at relevant spatial scales, e.g., the landscape-scale, and this is notoriously difficult. The gold standard in population abundance estimation is spatial-capture-recapture (SCR, Royle *et al*. 2014). The challenge is that SCR requires individually identifiable animals and requires detectors spaced relative to individual movement, with arrays large enough to expose a sufficient number of individuals to detection and detect the full extent of individuals movement, but close enough together that single individuals can be detected at multiple detectors (Sollmann et al. 2012; Royle et al. 2014; Sun et al. 2014). Meeting these competing requirements for multiple species with home ranges that vary over multiple orders of magnitude is often logistically implausible at the spatial scale at which management occurs, e.g., the area of a state or entire country. Consequently, SCR studies are often conducted at local scales and nevertheless remain underpowered with the inability to extrapolate to landscape-scales at which such policies are aimed, and wildlife population dynamics play out.

Site-structured models, for example, occupancy models (Mackenzie et al. 2002; Tyre et al. 2003) and N-mixture models (Royle, 2004), are hierarchical models that estimate the latent occupancy or abundance state of site *i* in the state model, and explicit model heterogeneity in detection probability conditional on the latent state in the observation model. Occupancy models leverage detection/non-detection data of unmarked animals from repeat surveys and can provide a logistically plausible and tractable alternative to abundance estimation. Occupancy models are best suited for low-density, wide-ranging species where site-level abundance does not vary, e.g., the number of individuals in an area is typically 0 or 1 such as observed in territorial carnivores, and thus occupancy will closely track the state variable of interest, abundance (Clare et al. 2015). For species which exhibit high levels of variation in site-level abundance, estimating occupancy can be a coarse and uninformative metric (z_i_ =1 for every site with at least one individual). Royle–Nichols models (Royle & Nichols 2003) instead assume that site-level detection probability increases with the number of individuals present, so that heterogeneity in detection frequency carries information about abundance; they can thereby estimate expected abundance and individual-level detection probability from detection/non-detection data alone (Royle & Nichols, 2003). While these models can be highly sensitive to assumption violations, empirical tests have shown they can recover spatial patterns in expected abundance comparable to spatial density surfaces from SCR under certain conditions, for example, as seen for fisher (*Pekania pennanti*), a low-density carnivore across a 70,000 km^2^ region of New York State (Linden *et al*. 2017). Royle-Nichols models have also been used to estimate expected abundance of difficult-to-monitor animals on landscape scales (Twining et al. 2023; 2024). The abundance parameter estimated is best interpreted as an index of abundance, as opposed to true abundance, with estimates being sensitive to assumption violations (Nakashima, 2020; Zwerts et al. 2026). While being cognizant of these caveats, occupancy and Royle-Nichols models provide capacity to provide inference on both distribution and abundance of wildlife populations while accounting for heterogeneity in detection from spatially extensive landscape-scale survey that would be otherwise infeasible (Gilbert et al. 2021)

The state of Mato Grosso in central-western Brazil provides a compelling case study to test the efficacy of the Forest Code in maintaining populations of medium and large-bodied mammals at a landscape-scale. Mato Grosso has been the epicenter of Brazil’s agricultural expansion for the past four decades and is now the country’s (and also one of the world’s), largest producers of cattle, soybean, and corn (Arvor et al. 2012; Spera et al. 2014). Alongside agricultural expansion, production has intensified through soybean–corn double cropping, in which soybean sown at the onset of the rains is followed by a second corn crop on the same fields, so that the same landscape cycles annually between two very different crop resources (Spera et al. 2014; Zalles et al. 2019). Despite being formed of three distinct biomes, the Amazon rainforest in the north, the Cerrado savanna in the center and east, and the Pantanal wetland in the south, the entire state of Mato-Grosso lies within the Legal Amazon, thus the Forest Code requirements are at their most stringent here (80% of native vegetation preserved in forested habitat, and 35% in Cerrado savanna habitat). The outcome is a mosaic in which large blocks of legally retained forest and savanna are embedded within a matrix of pasture and cropland (Stickler et al. 2013). Whether this mosaic supports populations of the state’s largest vertebrates, or merely their transient presence, is unknown.

Occupancy and abundance depend on biotic as well as abiotic factors, for predators, the availability of prey is critical. In Mato Grosso system, jaguars are the apex predator, with peccaries among their most important prey (Emmons 1987; Cavalcanti & Gese 2010; Hayward et al. 2016; Dymit et al. 2026). The depletion of ungulate prey bases has recently been implicated in jaguar declines (Ferraz et al. 2026). Ecological theory posits that the strength of a species interaction scales with the population size of the interacting species, thus prey abundance rather than prey presence is the relevant quantity to estimate, and recently developed abundance-mediated interaction models allow the occupancy of one species to be modelled as a function of the latent abundance of another from the same detection/non-detection data (Twining et al. 2025). In production landscapes where crops subsidize prey, such models can reveal whether prey abundance can compensate for limited native habitat.

Here, we use a landscape-scale camera trap survey conducted over a 10-month period, spanning both crop seasons, across a 606,000 km^2^ area of Mato-Grosso to quantify the occupancy and abundance of three medium- and large-bodied terrestrial vertebrates, jaguar, white-lipped peccary, and lowland tapir, across the state, and in doing so assess the conservation value of these multi-use production landscapes for these focal threatened species of conservation concern. We predicted that (1) jaguar occupancy would increase with forest cover and peccary abundance and decrease with agricultural land, distance from rivers, road density, and elevation; (2) peccary abundance would increase with forest cover and with soybean–corn cropland, a trophic subsidy, and decrease with distance from rivers and elevation; and (3) tapir abundance would increase with forest cover and decrease with cropland, elevation and distance from rivers.

## Methods

### Landscape-scale camera trap surveys

A landscape-scale camera trap survey was conducted across a 606,000 km^2^ region of the State of Mato Grosso in Brazil (see Figure 1), with sampling including both Amazonian and Cerrado biomes. During a 10-month period between November 2024 – Sept 2025, the 115 sites were sampled using camera traps for a mean of 71 days per site (range = 4 – 155 days). During this 10-month period there are two distinct crop growing seasons, and a third period of bare soil in the same spatial locations– from 1^st^ November to the end of February soybean is the primary crop grown, from while from 1^st^ March – 14 August, corn is grown in the same areas. Then between these seasons from 15 August to September is a post-harvest period, and the soil is left bare. Sites were spaced a minimum of 1.32 km apart distance apart with a mean pairwise distance of 30.85 km. All images were processed and animals identified to species. These were used to produce weekly detection histories for each species at each site.

**Figure 1.**
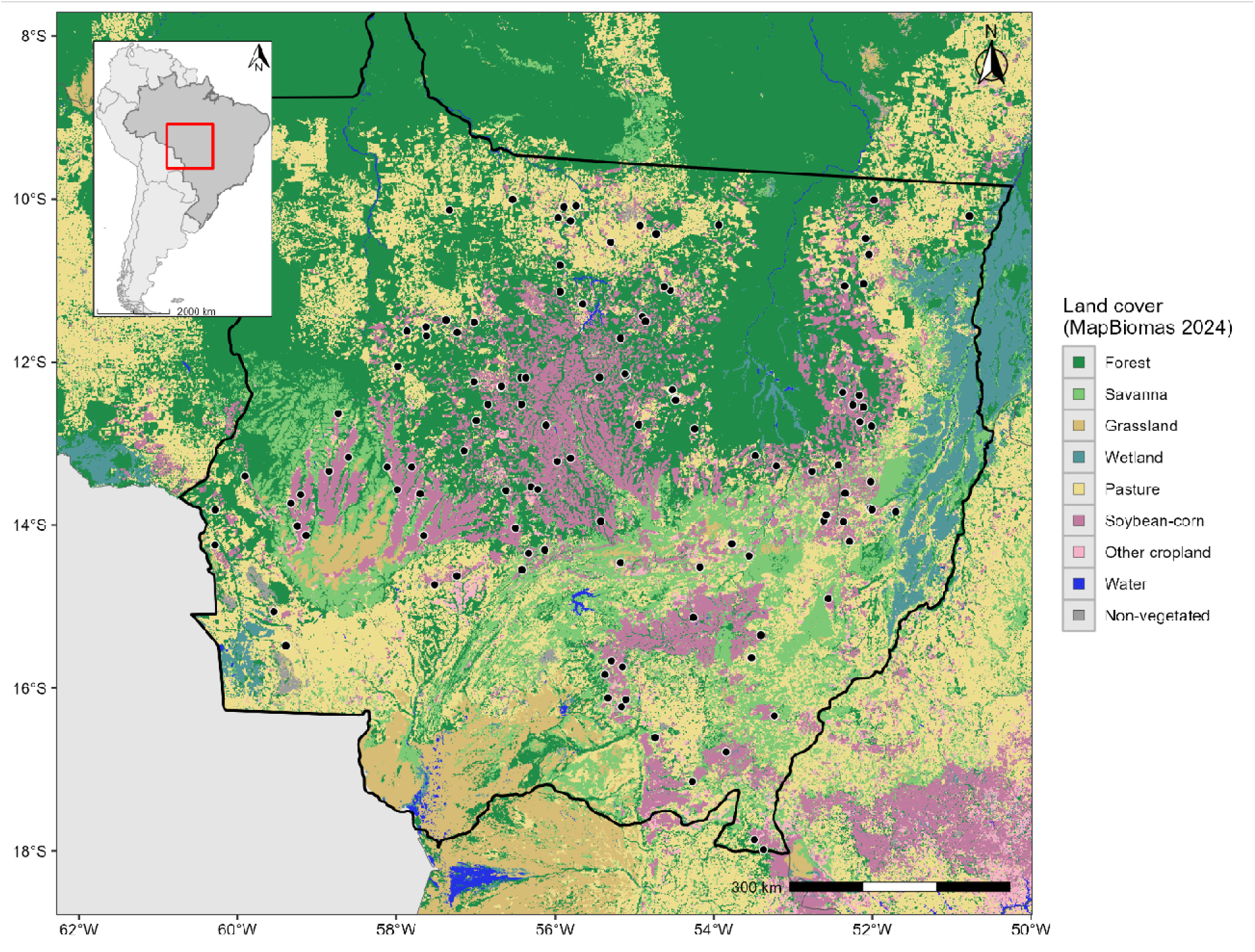
Map showing the 115 sites sampled (black circles) across the State of Mato-Grosso in Brazil. Land cover of the region is shown with habitats color-coded as described in legend. Note: the Xingu Indigenous Park (the large, forested area in the northeast of Mato-Grosso) was not sampled.

### Hierarchical modelling: single-species models

For jaguars we first fit a standard single-season occupancy model, where we assume our observations at site *i* on occasion *j* are the result of a draw from a Bernoulli distribution with probability *p_ij_* (detection probability) conditional on occupancy (z_i_ = 1):

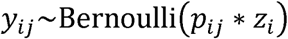

It is assumed that detection probability can be estimated as a function of site and site-by-occasion covariates using a logit-link:

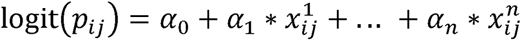

where *α_0_* is the intercept and α*_1_*… α*_n_* are slope terms for covariates *x^1^*… *x^n^*. Then in the state model the latent true occupancy state, *z_i_,* is estimated as a Bernoulli random variable:

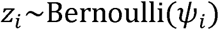

Where, *ψ_i_*, is the probability of occupancy at site *i,* and *z_i_*= 1 when a species occurs at site *i,* and *z_i_* = 0 when the species is absent from site *i.* It is assumed that the occupancy probability of a species at site *i* can be calculated as a function of site-level covariates using a logit-link:

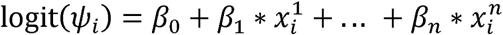

where *β*_0_ is the intercept (mean occupancy on the logit-scale), *β_n_* are slope terms, for associated site-level covariates *x_i_^n^*. For tapir and peccary, we fit single-season Royle-Nichols models (Royle & Nichols, 2003), these are an extension of occupancy models, also using detection/non-detection data, but exploiting an assumed link between site-level abundance and detection probability. Our observations at site *i* on occasion *j* (*y_ij_*) are now assumed to be a single draw (K=1) from a binomial distribution with the success parameter, *p_ij_*, representing population level detection probability at site I on occasion j:

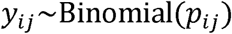

The population-level detection probability *p_ij_* is a function of both site-level abundance, N_i_, and individual detection probability, r_ij_, where p_ij_ is calculated as:

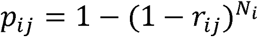

Where individual detection probability, r_ij_, can be calculated as a function of site-level covariates using a logit-link:

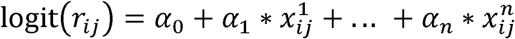

In the state model, site-level abundance (N_i_) is assumed to be a drawn from a Poisson distribution with mean and variance, *λ_i_*:

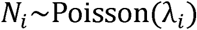

Where expected abundance at site i, *λ_i_*, can be calculated as a function of site-level covariates using a log-link:

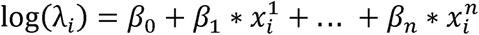

### Model fitting and model selection

Our global single-species models were species-specific and based on *a priori* hypotheses informed by literature and our knowledge of the study system. For each species we summarized covariates at spatial-scales relevant to their home range size in the Cerrado region of Brazil, where there was uncertainty, we used the lower end of published estimates. For jaguars this was 100km^2^ (Silveira, 2004), for white-lipped peccary this was 50 km^2^ (Costa et al. 2023), and for tapir, this was 5 km^2^ (Medici *et al*. 2022). All species also shared a set of candidate detection covariates including the effects of occasion number (time since deployment), effort (number of days the camera trap was functional during each 7-day occasion), and ordinal day (both linear and quadratic terms). Latitude and longitude were included as candidate covariates in the state model to account for broad spatial trends. In addition to the shared spatial and detection covariates, we hypothesized that jaguar occupancy would be positively associated with forest cover, and negative associated with pasture (prop.), distance from major rivers (m), road density (km/km^2^), and mean elevation (m). We hypothesized that white-lipped peccary abundance would be positively associated with crop-based trophic resources for which we consider land used for soybean and corn production, as well as forest cover, and negatively associated with distance to nearest river (m) and elevation (m). Finally, we hypothesized that tapir abundance would be positively associated with forest cover and negatively associated with agricultural land-uses, as well as elevation and distance to nearest river (m). Land cover proportions were calculated using the package ‘landscapemetrics’ (Hesselbarth et al. 2019) and MapBiomas, Brazil 2024 landcover maps. We calculated mean elevation using STRM-based elevation from AWS Terrain Tiles using the ‘elevatr’ package (Hollister et al. 2025). Road density was calculated using the Global Roads Inventory Project road-density rasters (Meijer et al. 2019), and distance to river was calculated using hydroRIVERS v1.0 (Lehner & Grill, 2013). To account for expected differences in focal species movement, occupancy, and abundance during the 10-month sampling period underpinned by the double cropping practices used on the landscape and the turnover of trophic resources on the landscape (1^st^ Nov – 28^th^ Feb = soybean, 1^st^ March – 14 August = corn, and 15^th^ August – October = bare soil), we used a stacked data design, whereby each site x season combination was considered a unique site. This resulted in 149 site x season combinations from 115 unique sites. There were only four site x season combinations that fell within the bare soil period, these were dropped due to insufficient data to support parameter estimation, leaving 145 sites for analysis. We fixed crop season (soybean = reference level, corn = slope term) on both the state and detection model across all models. All covariates were assessment for multi-collinearity using variance inflation factors (Zuur et al. 2010), with multi-collinearity defined as VIF values > 3. There was strong multi-collinearity between pasture (11.26), soybean/corn (11.95), and forest (7.77). We dropped pasture from the analysis, and then recalculated the VIF scores, after the removal of pasture there was no further evidence of collinearity among the retained covariates. All covariates are standardized to have a mean of 0 and SD of 1 to improve mathematical convergence and increase ease in interpretation of models’ coefficients.

All occupancy and Royle-Nichols models were fit using Maximum Likelihood in the package ‘unmarked’ (Kellner et al. 2023). We use Akaike’s Information Criterion (AIC) for identification of the most parsimonious model (Akaike, 1973; Burnham & Anderson, 2001; 2004). We used a two stage (secondary candidate set) model selection process (Morin *et al*. 2020). First, we conducted AIC-based model selection on the detection sub-model, with season fixed on *p,* considering all possible parameter combinations while keeping the state model as intercept only. Next, we fix the detection model as intercept only, and then we used AIC-based model selection to identify the best supported combination of parameters for the state model while keeping season fixed on ψ. Finally, we combined the top detection and top occupancy submodel to produce a final model used for each species. We identified redundant or ‘pretending’ parameters in both state and observation model by ensuring additional parameters improved model ranking by >2 AIC points (Arnold, 2010). Where a parameter is identified as redundant, the model is removed from consideration (we still show models containing redundant parameters in our AIC tables but highlight top models in bold [after dismissing models containing redundant parameters]). Season was retained in all models regardless of its support because the stacked data structure treats station × season combinations from the same station as independent. We provide all estimates from our top models with 85% confidence intervals to align with AIC model selection process (Sutherland *et al*. 2023).

### Hierarchical modelling: abundance-mediated interaction models

To test hypotheses on the interactions between habitat availability and prey abundance on jaguar occupancy in the landscape, we fit a two-species abundance mediated interaction model (Twining et al. 2025). These models use the same data and sub-models described above but allow the occupancy probability or expected abundance of one species to depend on the latent abundance of another species. Here, we built a two species model whereby the detection probability of jaguars at site *i* on occasion *j* was estimated as a function of the top site-by-occasion covariates selected above in the single species models (crop season only), and occupancy probability of jaguars at site *i* was estimated as a function of top habitat covariates selected (forest, elevation, and crop season (soybean vs. corn), as well as latent peccary abundance at site *i*. We specified a Royle-Nichols model for peccary to estimate latent abundance at site *i* using the covariates from the top single species model (soybean/corn and forest). We modified the linear predictors for jaguar occupancy probability to represent the hypothesis that jaguar occupancy is positively associated with peccary abundance:

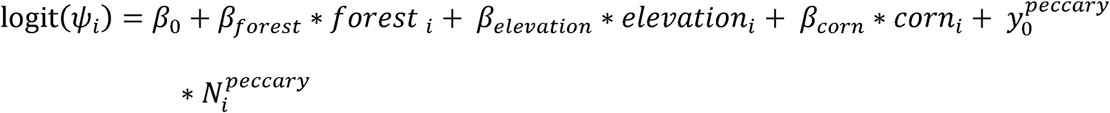

Where *y*_0_*^peccary^* is the intercept for a constant interaction term between peccary abundance and jaguar occupancy probability, and *N_i_^peccary^* is the latent abundance of peccary at site *i.* We developed a new function ‘occuRNMulti’ in package ‘unmarked’ (Kellner et al. 2023) to fit abundance-mediated interaction models using maximum likelihood (Code S1). We report parameter coefficients and present conditional effects plots with 85% confidence intervals consistent with the AIC-based model selection criteria used in the first stage.

## Results

### Single species models

#### Jaguar

Jaguars were detected at 45 of the 145 site x season combinations, with a total of 91 weekly detections. The final model was for jaguar included detection probability being modelled as a function of season (which was fixed, with none of the other covariates explaining variation in detection probability), and occupancy probability being modelled as a function of season, forest, and elevation (see Table S1 and S2). Jaguar mean weekly detection probability was 0.15 (85% CI = 0.12, 0.18), with the season not explaining significant variation in detection probability y (α_corn_ = 0.21, 85% CI = -0.43, 0.85). The top model estimated an occupancy probability for jaguars during the soybean season at mean covariate values of 0.45 (85% CI = 0.33 – 0.57). The top model for explaining variance in occupancy probability included forest (β_forest_ = 0.99, 85% CI = 0.47, 1.52) and elevation ( β_elevation_= -0.91, 85% CI = -1.39, -0.41), along with the fixed effect of season (β_corn_ = -2.13, 85% CI = -3.48, -0.77, see Table S2).

Conditional effects plots reveal that at mean elevation (415 m), an increase in forest cover from 15% of a 100 km^2^ grid cell to 65% increased the probability of jaguar occupancy from 0.33 (85% CI = 0.23 – 0.44) to 0.87 (85% CI = 0.64 – 0.97, see Figure 2A). While at mean forest cover (0.24), the probability of jaguar occupancy dropped from 0.78 (85% CI = 0.59 – 0.89) at 200 m above sea level to 0.05 (85% CI = 0.01 – 0.22) at 800 m above sea level. Finally, occupancy probability for jaguars was far lower during the corn growing season than soybean season, dropping from 0.45 (85% CI = 0.33 – 0.57) during the soybean season, to 0.09 (85% CI = 0.03 – 0.24) during the growing season for corn. See conditional effects plots in Figure 2A for visualization of effect sizes on the natural scale. See Figure 3A for expected abundance predictions on a landscape-scale as a function of the interactions between forest cover and elevation.

**Figure 2.**
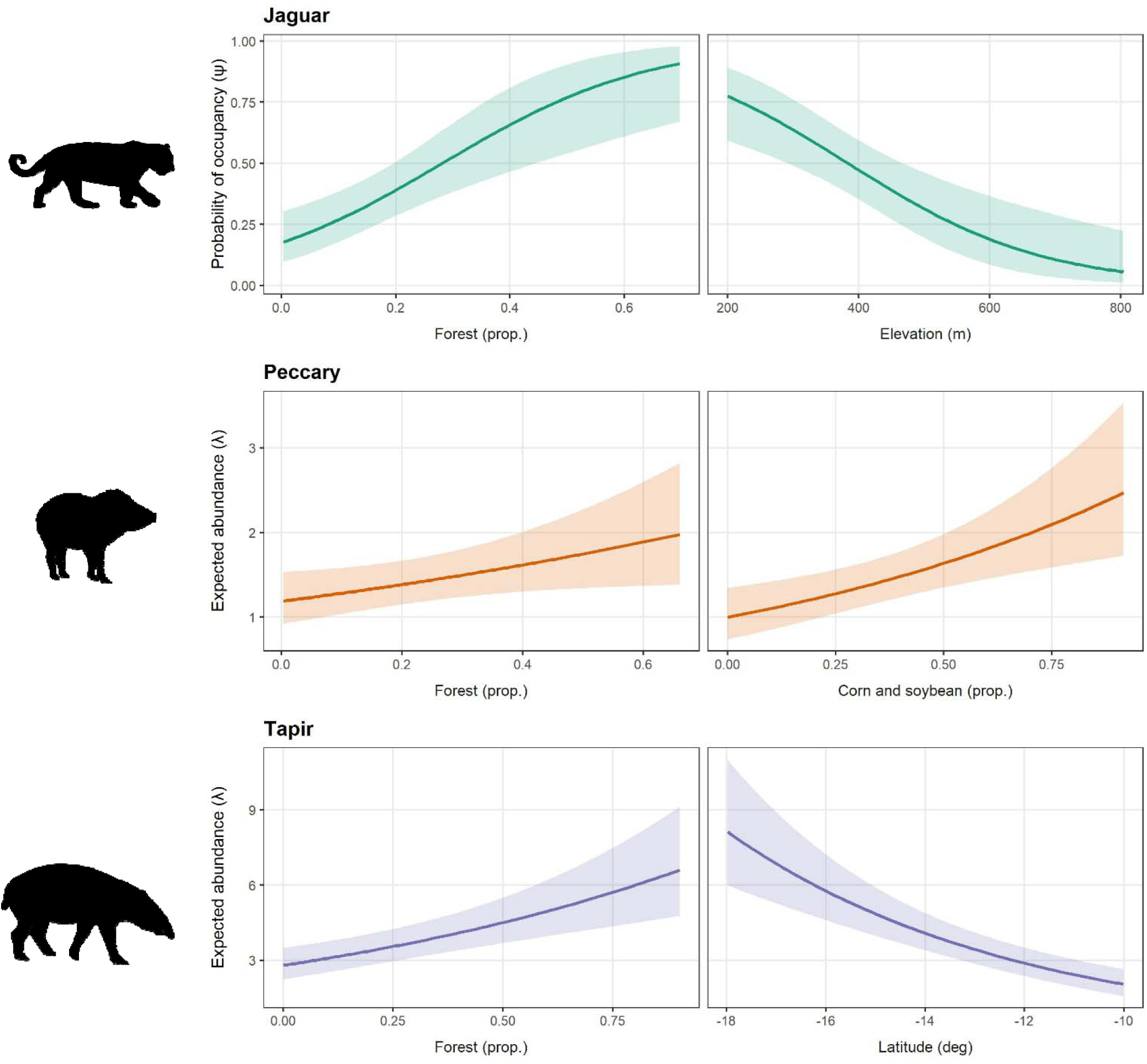
Conditional effects plots from the top-ranked model for jaguar (top), white-lipped peccary (middle), and tapir (bottom) model showing: A) predicted probability of occupancy of jaguars as a function of proportion of a 100 km^2^ grid composed of forest and elevation above sea level (m), B) expected abundance of white-lipped peccary as a function of proportion of a 50 km^2^ grid cell composed of forest and corn/soybean, and C) expected abundance of tapir as a function of a 5 km^2^ grid cell composed of forest and as well as latitude (decimal degrees). Predictions are for the soybean season with other covariates held at their means; shaded bands are 85% CIs.

**Figure 3.**
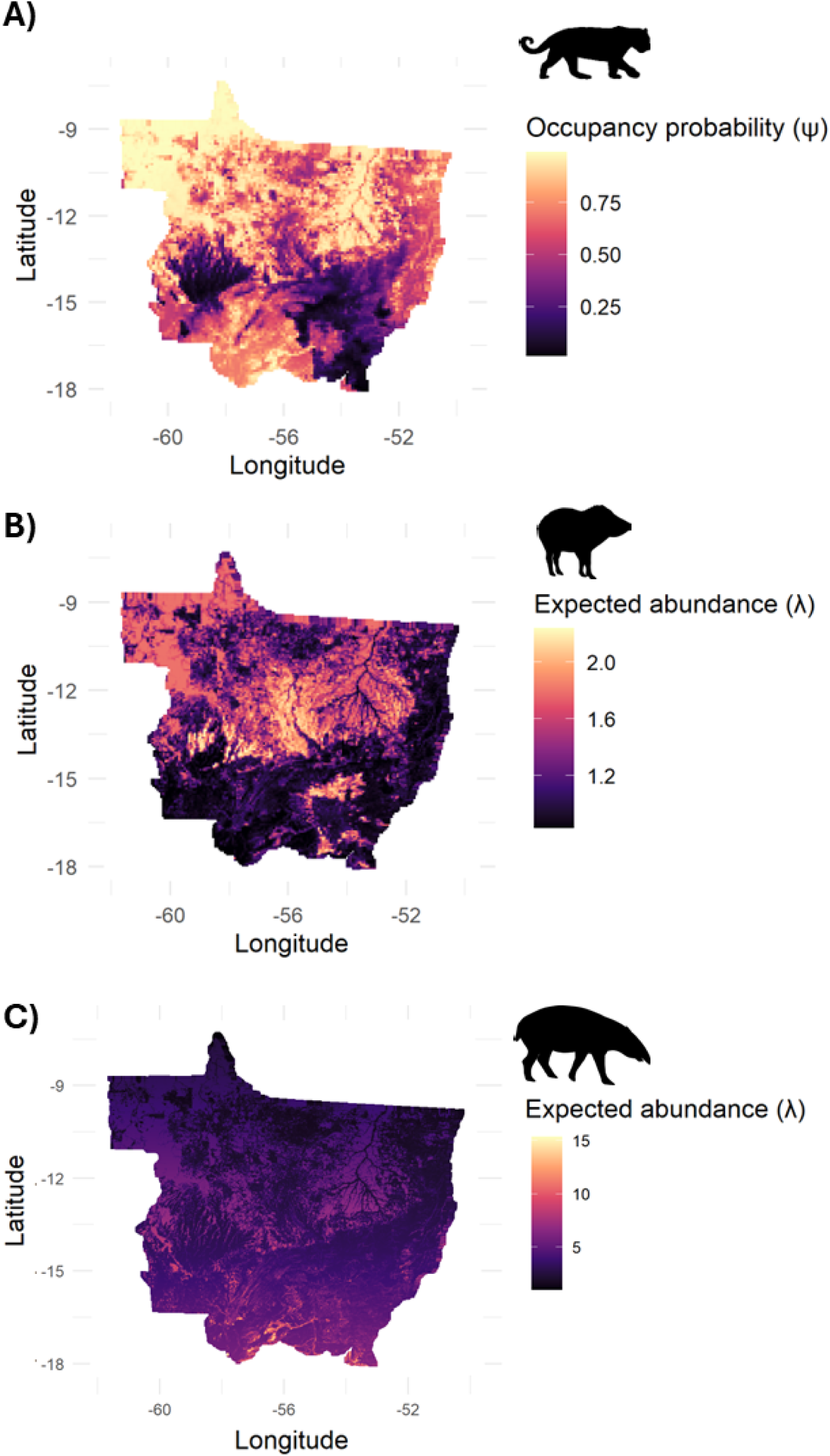
Spatially explicit predictions across the state of Mato Grosso, Brazil, from the top-ranked model for jaguar (top), white-lipped peccary (middle), and tapir (bottom) model showing: **A)** predicted probability of occupancy of jaguars as a function of proportion of a 100 km^2^ grid composed of forest and elevation above sea level (m), **B)** expected abundance of white-lipped peccary as a function of forest and corn/soybean cover, and **C)** expected abundance of tapir as a function of forest and latitude (decimal degrees).

#### White-lipped peccary

Peccary were detected at 93 of the 145 site x season combinations, with a total of 330 weekly detections. The final model for peccary included crop season (α_corn_ = -1.27, 85% CI = -2.22, -0.32), effort (α_effort_= 0.22, 85% CI = 0.10, 0.33), and ordinal date (α_date_ = 0.32, 85% CI = 0.22, 0.41, see Table S3) on detection probability, with time (occasion) and the quadratic form of ordinal date being redundant parameters (see Table S3). Peccaries mean weekly individual-level detection probability during the soybean season was 0.21 (85% CI = 0.18, 0.24). The final model for expected abundance of peccary included forest (β_forest_= 0.14, 85% CI = 0.01, 0.27) and soybean/corn production (β_soybean/corn_ = 0.21, 85% CI = 0.08, 0.34), along with the fixed effect of season (β_corn_ = 0.58, 85% CI = -0.26, 1.43, see Table S4). The final model estimated a mean expected abundance for peccary of 1.42 (85% CI = 1.18 – 1.71)

Model estimates demonstrate that expected abundance of peccary increases by 15% (85% CI = 1% increase, 31% increase) per SD unit increase in forest cover at a 50 km^2^ scale. Similarly expected abundance of peccary increases by 23% (85% CI = 8% increase, 40% increase) for every SD unit increase of soybean and corn cover in a 50 km^2^ area. There is a general trend for peccary abundance to be higher during the corn growing season, with expected abundance on average 79% higher than during the soybean growing season, however, uncertainty was extremely high and confidence intervals overlapped zero (85% CI = 23% decline, 318% increase). See conditional effects plots in Figure 2B for visualization of effect sizes on the natural scale. See Figure 3B for expected abundance predictions on a landscape-scale as a function of the interactions between forest cover and latitude.

#### Lowland tapir

Tapir were detected at 135 of the 145 site x season combinations, with a total of 871 weekly detections. The final model for tapir estimated a mean weekly individual detection probability during the soybean season of 0.29 (85% CI = 0.24, 0.33). Model selection revealed time (occasion) and both the linear and quadratic form of ordinal date to be redundant parameters. The final model included the fixed effect of season (α_corn_= 0.45, 85% CI = 0.0, 0.9), along with effort (α_effort_ = 0.32, 85% CI = 0.24, 0.39, see Table 5), the latter clearly and the former marginally positively associated with detection probability. The final model estimated a mean expected abundance for tapir of 3.6 (85% CI = 3.05, 4.36). The top-ranked model revealed a positive association between expected tapir abundance and forest cover (β_forest_ = 0.21, 85% CI = 0.11, 0.31) and a negative association with latitude (β_latitude_= -0.31, 85% CI = -0.41, -0.21), along with the fixed effect of season (β_corn_ = -0.19, 85% CI = -0.54, 0.16, see Table S6).

Model estimates demonstrate that expected abundance of lowland tapir increases by 23% (85% CI = 12% increase, 36% increase) per SD unit increase in forest cover at the 5 km^2^ scale. Expected abundance of tapir however decreased by 27% (85% CI = 19% decrease, 34% decrease) for every SD unit increase of latitude (i.e., from south to north). There was a slight trend for expected abundance of tapir to be lower during the corn growing season, with expected abundance on average 18% lower than during the soybean growing season, however, the confidence intervals overlapped zero (85% CI = 42% decline, 17% increase). See conditional effects plots in Figure 2C for visualization of effect sizes on the natural scale. See Figure 3C for expected abundance predictions on a landscape-scale as a function of the interactions between forest cover and latitude.

#### Abundance-mediated interaction models

Our abundance-mediated interaction model provides evidence of a positive association between jaguar occupancy probability and peccary abundance (γ_0_ = 0.50, 85% CI = 0.08 – 0.91). Jaguar occupancy remained positively associated with forest (□_forest_ = 0.97, 85% CI = 0.40, 1.54), and negatively associated with elevation (□_elevation_ = -1.07, 85% CI = -1.65, -0.49). Conditional estimates of occupancy conditional on low peccary abundance (N = 0) and high peccary abundance (N = 3) along the observed gradient of forest cover illustrate the joint effects of habitat and prey abundance on jaguar occurrence (Figure 4). At low peccary abundance (N = 0) and low forest cover (10% of 100 km^2^ cell), mean jaguar occupancy probability was 0.16 (85 CI% 0.07 – 0.30), but at the same low forest cover (10%), with high peccary abundance (N = 3), mean jaguar occupancy probability was almost than threefold (x2.9) higher at 0.45 (85% CI = 0.24 – 0.66). At high forest cover (65% of 100 km^2^ cell), the very large effect of peccary abundance on probability of jaguar occupancy diminishes somewhat, at low peccary abundance, mean jaguar occupancy probability was 0.78 (85% CI = 0.41 – 0.94), which increases more marginally at high peccary abundance to 0.94 (85% CI = 0.71 – 0.99).

**Figure 4.**
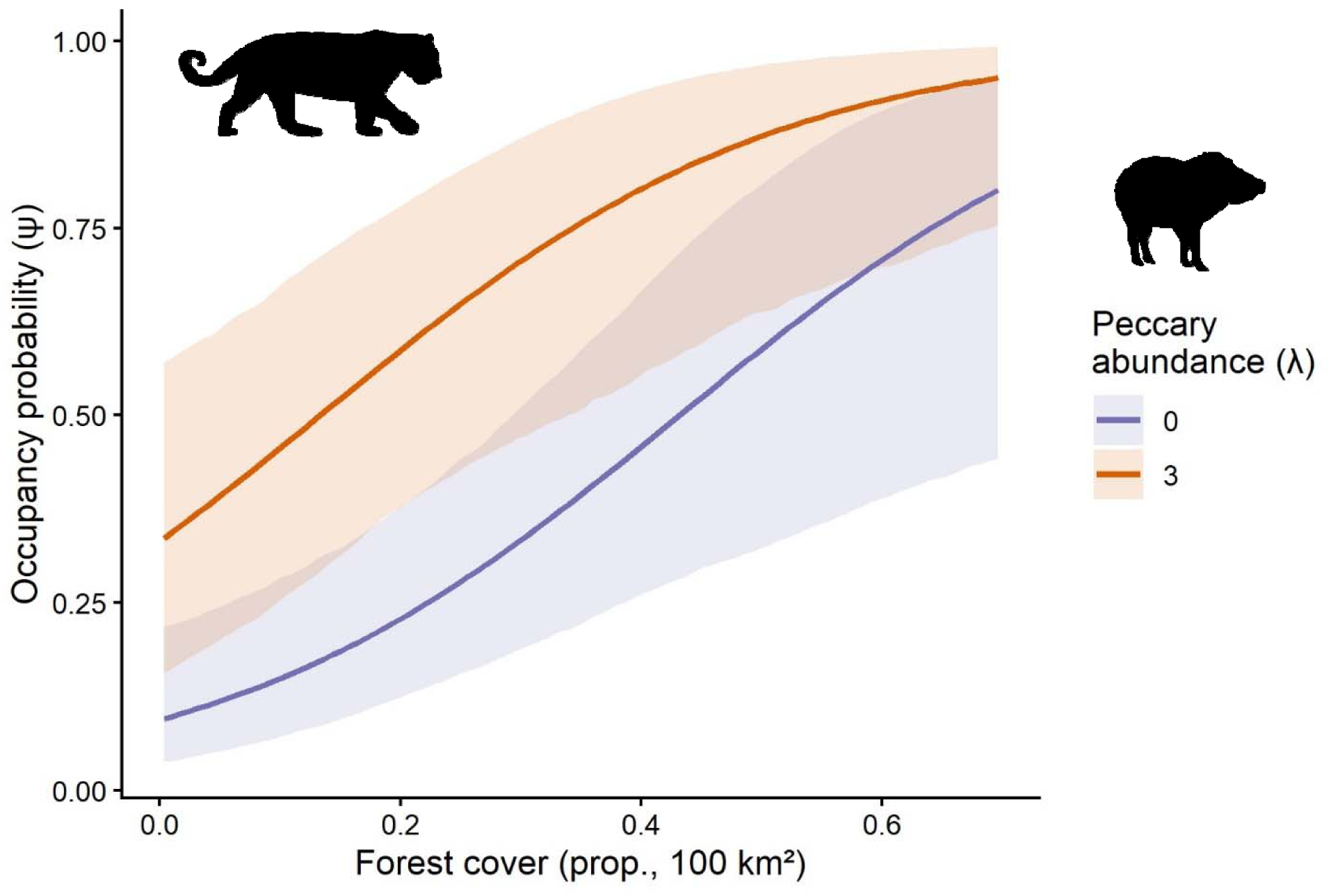
Predicted probability of jaguar occupancy across a gradient of forest cover conditional on low (N=0) and high (N=3) site-level peccary abundance. Season was set to soybean and all other covariates were kept at their mean. Shaded bands are 85% CIs.

## Discussion

Here, we present spatially explicit estimates of probability of occupancy and expected abundance of three threatened large-bodied mammals across ∼600,000 km^2^ in Mato-Grosso, Brazil, one of the world’s largest tropical agricultural production landscapes. By modelling detection/non-detection data from a landscape-scale camera trap survey with hierarchical models that account for heterogeneity in detection, we provide the first assessment of whether native vegetation retained on private lands under Brazil’s Forest Code sustains populations of large vertebrates, at the scale at which the policy operates. The empirical evidence is unambiguous, all three species were widespread, with derived occupancy estimates varying from 65 – 99%, demonstrating the high value of the Forest Code provisions in conserving populations of large-bodied mammals within a globally important production landscape. Forest cover was the only covariate retained in the top model for every species, and its effect was consistent across species and state variables: jaguar occupancy increased from 0.33 to 0.87 as forest cover in a 100 km^2^ cell increased from 15% to 65%, while expected abundance of tapirs and peccaries increased by 23% and 15% respectively, per standard deviation increase in forest cover. Our sampling was conducted within the private production landscapes of Mato Grosso, the native forest that remains is largely retained due to federal regulations, and the results provide direct evidence that the Forest Code sustains not only the occupancy but also local abundance of both an apex predator and two large-bodied herbivores within one of the mostly intensely farmed regions on earth.

Our results extend what previous studies have shown in individual forest fragments and corridors (at the scale of tens to hundreds of km^2^, Lees & Peres, 2008; Zimbres et al. 2017) to a landscape-scale. Comparing our occupancy and expected abundance estimates with existing estimates from across south and central America reveal both areas of consensus and incongruence. Our mean jaguar occupancy during the soybean season was 0.45 (85% CI = 0.33 – 0.57), which sits comfortably in the range of reported occupancy estimates from multi-site camera trap surveys across central America (where mean jaguar occupancy ranged from = 0.19 – 0.79, Calderón et al. 2022). In contrast, the near ubiquitous occurrence of tapirs observed in our survey across Mato Grosso exceeds existing estimates from the Orinoquia region, a comparable savanna-dominated production landscape in Colombia (mean ψ = 0.4 – 0.6, Alviz et al. 2025). While white-lipped peccary have been extirpated from approximately 87% of their historical range in Mesoamerica (Thornton et al. 2020), and one fifth of their total historical range (Altricher et al. 2012), the species remains abundant across the production landscapes of Mato Grosso, consistent with the ‘paradoxical’ persistence of the species previously reported for the state (Lima et al. 2019). This is likely linked to the forest reserves maintained in the landscape, and the species positive association with forest cover, as seen elsewhere e.g., in Paraguay (Weiler et al. 2020). The apparent success of the Forest Code in Mato Grosso seen here should however be interpreted in its regional context. The production landscapes and private forest reserves of Mato Grosso are bordered by very large continuous blocks of native habitat, the largest being the Amazonian forest to the north and the Xingu Indigenous Park to the northeast (see Figure 1). These two protected areas may drive continuous source-sink dynamics that allow herd forming peccaries to recolonize after regional population crashes characteristic of the species (Fragoso et al. 2022; Painkow Neto et al. 2026), while supporting abundant tapir populations, and jaguars. The conservation value of retained habitat within production landscapes is therefore best viewed as complementary to, rather than as a substitute for large, protected areas and Indigenous territories.

Jaguar occupancy in the landscape was primarily underpinned by variation in forest cover, elevation, and peccary abundance, and the interactions amongst these variables. The strong positive association of jaguar occupancy with forest cover is consistent with a range-wide analysis showing that jaguars select for productive forested habitat (Jędrzejewski et al. 2018; Portugal et al. 2020; Alvarenga et al. 2025). The negative association with elevation most plausibly reflects the geography of Mato Grosso as opposed to a simple elevational effect: the higher plateaus are the oldest and most intensively managed soybean and corn districts, whereas the low-lying north and the river valleys retain more forest with higher primary productivity, which is the strongest predictor of jaguar density across the species’ range (Jędrzejewski et al. 2018).

Our abundance-mediated interaction model provides evidence that jaguar occurrence is associated with prey as well as habitat: jaguar occupancy increased with latent peccary abundance (γ□= 0.50), and the effect on the probability scale was greatest where forest was scarce. At 10% forest cover, occupancy almost tripled from where peccaries were absent to where they were abundant (0.16→0.45), whereas at 65% forest cover, jaguar occupancy was already high and rose more modestly with increasing peccary abundance (0.78→0.96). Peccaries, and white-lipped peccaries in particular, are among the most frequently taken prey of jaguars across their Neotropical range, although jaguars diet displays pronounced variation (Emmons 1987; Polisar et al. 2003; Cavalcanti & Gese 2010; Sollmann et al. 2013; Hayward et al. 2016; Foster & Harmsen 2022; Dymit et al. 2026). The loss of the ungulate prey base has been identified as a principal driver of jaguar decline in the Atlantic Forest (de Barros Ferraz et al. 2026). Modelling the interaction between jaguar and peccary as a function of abundance rather than co-occurrence matters because interaction strength scales with population size, and such signals can be lost for widespread species in occupancy-only formulations (Twining et al. 2025). Our results show that agricultural production and jaguar presence are not necessarily mutually exclusive provided that sufficient native forest is maintained across the landscape. A practical implication is that jaguar persistence in these landscapes may depend not only on how much forest is retained but also on prey availability: where forest is limited, abundant peccaries, subsidized by crops, may still allow jaguars to persist, although jaguars drawn into low-forest, peccary-rich crop districts are also likely to face a higher risk of retaliatory killing (Michalski et al. 2006), with the potential to act as an ecological trap.

The two herbivores differed in their responses to the habitats within the production landscape. The expected abundance of white-lipped peccary was positively associated with forest cover and extent of soybean and corn cropland, which indicates the population in Mato Grosso depend on a combination of native forest and crop resources rather than either alone. GPS data from the same region and from the Cerrado show this same pattern (Costa et al. 2023; Painkow Neto et al. 2024). These results align with the observation that crop-raiding risk across Mato Grosso is highest precisely where forest cover and connectivity are greatest (Costa et al. 2026). The distinction between the two crops observed in our parameter estimates is ecologically important: peccaries feed intensively on second-crop corn but only occasionally on soybean (Lima et al. 2019; Painkow Neto et al. 2024). Because our cropland covariate pooled the two crops, the analysis can separate them only through season; the tendency for expected abundance to be higher during the corn season (79%) is consistent with this distinction but was estimated with wide uncertainty, and the crop × season interaction was not supported (Table S4). Lowland tapirs were the most frequently detected of the three species, occurring at almost every site. Unlike peccaries, their expected abundance was not associated with cropland and instead was positively associated with forest cover, and negatively with latitude. Their widespread abundance in the landscape may in part by explained by their foraging ecology; as browsers with a broad diet including leaves, shoots, fruits and seeds (Fragoso, 1991; Fragoso & Huffman, 2000; Henry et al. 2000; O’Farrill et al. 2013), tapirs can exploit resources throughout the agricultural cycle, and they make intensive use of early-successional forest (de Luca & Pardini 2017), which is abundant in the mosaic of forest remnants and fields in Mato Grosso. Intriguingly, in our analysis, tapir abundance was observed to decline with increasing latitude, that is, from the southern Cerrado and transitional zone towards the Amazonian north. This latitudinal gradient likely captures variation in climate, productivity, biome composition, land-conversion history, and hunting pressure that our other covariates did not. This result stands in contradiction to our expectation that large mammals should necessarily be most abundant in the least converted, most Amazonian portions of the state, and suggests that long-established production landscapes that have retained forest reserves can support tapir occupancy. The high expected abundance in production landscapes observed here may, however, mask another ecological trap: telemetry shows that tapir movement is insensitive to human disturbance (Medici et al. 2022), so tapirs in these landscapes are exposed to roads, dogs, hunting and non-target poisoning (Lima et al. 2019), and a 10-month snapshot cannot distinguish self-sustaining populations from those maintained by immigration.

While our data is observational, and thus the evidence produced is correlational in nature, we apply probabilistic hierarchical models that explicitly account for heterogeneity in detection using a representative sample from across an entire production landscape. Our data is from a single year, the seasonal effects we report confound crop phenology with climatic seasonality, and a 10-month detection/non-detection survey cannot distinguish self-sustaining populations from those maintained by immigration from the Amazon and the Xingu. Nonetheless, abundance-mediated interaction models applied here provided critical insights into the factors that drive the occurrence of our focal species that were otherwise not visible. Critically, the model estimates demonstrate that jaguar occurrence is not underpinned by habitat and forest cover alone, but as has been suggested elsewhere, depends on the local abundance of primary prey species such as peccary. It is important to recognize that multi-species models like the abundance-mediated interaction model applied here are data-hungry: 145 site x season combinations are at the lower limit for reliable inference on interaction parameters with moderate effect sizes as seen here (Cowans et al. 2025; Twining et al. 2025; Twining & Kellner 2026). Additionally, it’s important to recognize that a positive association between jaguar occupancy and peccary abundance could arise from unmodelled heterogeneity in the state process, e.g., shared habitat preferences not captured by the habitat and spatial covariates in the model; we therefore present this result as consistent with, rather than proof of, prey-mediated jaguar occurrence. Finally, we had no covariates to serve as a proxy for hunting pressure or retaliatory killing which are the mechanisms most likely to convert the high occupancy and abundance we document into population sinks, and thus the scale of such actions, and potential to drive decline the region require further examination.

In conclusion, our results demonstrate that the native forest retained on private land under Brazil’s Forest Code in Mato Grosso is associated with the persistence and local abundance an apex predator, a megaherbivore and a keystone ungulate across an entire production region, and challenges the perception that landscapes dominated by mechanized agriculture are necessarily empty of large vertebrates. The Legal Reserves maintained under federal regulation appear to be functioning as suitable habitat for threatened species in Mato Grosso, although a more direct test would require comparing properties by compliance status and including reference sites in protected and / or Indigenous lands. Because continued persistence of large vertebrates depends on tolerance, habitat-retention policy should be paired with the mitigation of crop damage and livestock depredation, to reduce conflict between the animals these forest reserves sustain and the landholders who host them. The analytical framework used here, combining occupancy, Royle–Nichols and abundance-mediated interaction models fitted to a single landscape-scale camera-trap design, offers a transferable template for evaluating habitat-retention policies for wide-ranging, difficult-to-monitor species elsewhere.

## Data accessibility statement

All data, analyses code, and new ‘unmarked’ package functionality is hosted on a private anonymous Zenodo repository (https://zenodo.org/records/23109008). The files can be accessed by reviewers using this link: https://shorturl.at/THWNP.

