## Supplemental Materials for "Private forest reserves in Brazil’s agricultural landscapes effectively sustain the occupancy and abundance of multiple threatened large vertebrates"

**Data and code accessibility statement**: All data, analyses code, and new ‘unmarked’ package functionality is hosted on a private anonymous Zenodo repository (<https://zenodo.org/records/23109008>). The files can be accessed by reviewers using this link: <https://shorturl.at/THWNP>.

**Table S1.** Model selection table for the jaguar detection sub-model where models are ranked by AIC from lowest to highest. Table includes observation model specification, number of parameters (K), the log-likelihood (ℓ), and both AIC (2K - 2ℓ), and ΔAIC (AIC – lowest AIC). The top model after consideration of redundant parameters is highlighted in bold.

| **Observation model** | **Number of parameters (*K*)** | **Log-likelihood (ℓ)** | **AIC** | **ΔAIC** |
| --- | --- | --- | --- | --- |
| **α_0_ + α_1_ * season_i_** | **3** | **-310.54** | **627.07** | **0.00** |
| α_0_ + α_1_ * season_i_ + α_2_ * effort_i,j_ | 4 | -310.17 | 628.34 | 1.27 |
| α_0_ + α_1_ * season_i_ + α_2_ * date_i,j_ | 4 | -310.45 | 628.89 | 1.82 |
| α_0_ + α_1_ * season_i_ + α_2_ * occasion_i,j_ | 4 | -310.49 | 628.97 | 1.90 |
| α_0_ + α_1_ * season_i_ + α_2_ * date^2^_i,j_ | 4 | -310.50 | 629.01 | 1.94 |

**Table 2.** Model selection table for the jaguar occupancy sub-model where models are ranked by AIC from lowest to highest. Table includes state model specification, number of parameters (K), the log-likelihood (ℓ), and both AIC (2K - 2ℓ), and ΔAIC (AIC – lowest AIC). The top model after consideration of redundant parameters is highlighted in bold.

| **State model** | **Number of parameters (*K*)** | **Log-likelihood (ℓ)** | **AIC** | **ΔAIC** |
| --- | --- | --- | --- | --- |
| **β_0_ + β_1_ * season_i_ + β_2_ * forest_i_ + β_3_ * elevation_i_** | **5** | **-293.99** | **597.98** | **0.00** |
| β_0_ + β_1_ * season_i_ + β_2_ * forest_i_ + β_3_ * elevation_i_ + β_4_ * river distance_i_ | 6 | -293.88 | 599.75 | 1.77 |
| β_0_ + β_1_ * season_i_ + β_2_ * forest_i_ + β_3_ * elevation_i_ + β_4_ * longitude_i_ | 6 | -293.94 | 599.88 | 1.90 |
| β_0_ + β_1_ * season_i_ + β_2_ * forest _+_ β_3_ * elevation + β_4_ * latitude_i_ | 6 | -293.97 | 599.93 | 1.95 |
| β_0_ + β_1_ * season_i_ + β_2_ * forest_i_ + β_3_ * elevation_i_ + β_4_ * road density_i_ | 6 | -293.99 | 599.97 | 1.99 |

**Table 3.** Model selection table for the white-lipped peccary observation sub-model where models are ranked by AIC from lowest to highest. Table includes observation model specification, number of parameters (K), the log-likelihood (ℓ), and both AIC (2K - 2ℓ), and ΔAIC (AIC – lowest AIC). The top-ranked model after consideration of redundant parameters is highlighted in bold.

| **Observation model** | **Number of parameters (*K*)** | **Log-likelihood (ℓ)** | **AIC** | **ΔAIC** |
| --- | --- | --- | --- | --- |
| α_0_ + α_1_ * season_i_ + α_2_ * effort_i,j_ + α_3_ * occasion_i,j_ + α_4_ * date_i,j_ + α_5_ * date^2^_i ,j_ | 7 | -635.06 | 1284.12 | 0.00 |
| α_0_ + α_1_ * season_i_ + α_2_ * effort_i,j_ + + α_3_ * date_i,j_ + α_4_ * date^2^_i ,j_ | 6 | -636.18 | 1284.37 | 0.24 |
| α_0_ + α_1_ * season_i_ + α_2_ * effort_i,j_ + α_3_ * occasion_i,j_ + α_4_ * date_i,j_ | 6 | -636.47 | 1284.93 | 0.81 |
| α_0_ + α_1_ * season_i_ + α_2_ * effort_i,j_ + α_3_ * occasion_i,j_ + α_4_ * date^2^_i ,j_ | 6 | -636.95 | 1285.90 | 1.78 |
| **α_0_ + α_1_ * season_i_ + α_2_ * effort_i,j_ + α_3_ * date_i,j_** | **5** | **-637.99** | **1285.97** | **1.85** |

**Table 4.** Model selection table for the white-lipped peccary for abundance sub-model where models are ranked by AIC from lowest to highest. Table includes state model specification, number of parameters (K), the log-likelihood (ℓ), and both AIC (2K - 2ℓ), and ΔAIC (AIC – lowest AIC). The top-ranked model after consideration of redundant parameters is highlighted in bold.

| **State model** | **Number of parameters (*K*)** | **Log-likelihood (ℓ)** | **AIC** | **ΔAIC** |
| --- | --- | --- | --- | --- |
| β_0_ + β_1_ * season_i_ + β_2_ * forest_i_ + β_3_ * soybean/corn_i_ + β_4_ * longitude_i_ | 6 | -656.99 | 1325.99 | 0.00 |
| β_0_ + β_1_ * season_i_ + β_2_ * forest_i_ + β_3_ * soybean/corn_i_ + β_4_ * longitude_i_ + β_5_* river distance_i_ | 7 | -656.04 | 1326.07 | 0.08 |
| **β_0_ + β_1_ * season_i_ + β_2_ * forest_i_ + β_3_ * soybean/corn_i_** | **5** | **-658.07** | **1326.13** | **0.15** |
| β_0_ + β_1_ * season_i_ + β_2_ * forest_i_ + β_3_ * soybean/corn_i_ + β_4_ * river distance_i_ | 6 | -657.38 | 1326.76 | 0.77 |
| β_0_ + β_1_ * season_i_ + β_2_ * forest_i_ + β_3_ * soybean/corn_i_ + β_4_ * longitude_i_ + β_5_ * soybean/corn:season_i_ | 7 | -656.38 | 1326.77 | 0.78 |
| β_0_ + β_1_ * season_i_ + β_2_ * forest_i_ + β_3_ * soybean/corn_i_ + β_4_ * longitude_i_ + β_5_* river distance_i_ + β_6_ * soybean/corn:season_i_ | 8 | -655.54 | 1327.09 | 1.10 |
| β_0_ + β_1_ * season_i_ + β_2_ * forest_i_ + β_3_ * soybean/corn_i_ + β_4_ * soybean/corn:season_i_ | 6 | -657.67 | 1327.33 | 1.34 |
| β_0_ + β_1_ * season_i_ + β_2_ * forest_i_ + β_3_ * soybean/corn_i_ + β_4_ * longitude_i_ + β_5_* river distance_i_ + β_6_ * elevation_i_ | 8 | -655.71 | 1327.41 | 1.42 |
| β_0_ + β_1_ * season_i_ + β_2_ * forest_i_ + β_3_ * soybean/corn_i_ + β_4_ * longitude_i_ + β_5_* latitude_i_ + β_6_ * river distance_i_ _i_ | 8 | -655.80 | 1327.60 | 1.61 |
| β_0_ + β_1_ * season_i_ + β_2_ * forest_i_ + β_3_ * soybean/corn_i_ + β_4_ * river distance_i_ + β_5_ * elevation_i_ | 7 | -656.87 | 1327.75 | 1.76 |
| β_0_ + β_1_ * season_i_ + β_2_ * forest_i_ + β_3_ * soybean/corn_i_ + β_4_ * longitude_i_ + β_5_* latitude_i_ | 7 | -656.92 | 1327.84 | 1.85 |
| β_0_ + β_1_ * season_i_ + β_2_ * forest_i_ + β_3_ * soybean/corn_i_ + β_4_ * latitude_i_ | 6 | -657.96 | 1327.92 | 1.93 |
| β_0_ + β_1_ * season_i_ + β_2_ * forest_i_ + β_3_ * soybean/corn_i_ + β_4_ * longitude_i_ + β_5_ * elevation_i_ | 7 | -656.98 | 1327.97 | 1.98 |

**Table 5.** Model selection table for the lowland tapir observation sub -model where models are ranked by AIC from lowest to highest. Table includes observation model specification, number of parameters (K), the log-likelihood (ℓ), and both AIC (2K - 2ℓ), and ΔAIC (AIC – lowest AIC). The top-ranked model after consideration of redundant parameters is highlighted in bold.

| **Observation model** | **Number of parameters (*K*)** | **Log-likelihood (ℓ)** | **AIC** | **ΔAIC** |
| --- | --- | --- | --- | --- |
| α_0_ + α_1_ * season_i_ + α_2_ * effort_i,j_ + α_3_ * occasion_i,j_ + α_4_ * date_i,j_ + α_5_ * date^2^_i ,j_ | 7 | -734.05 | 1482.10 | 0.00 |
| α_0_ + α_1_ * season_i_ + α_2_ * effort_i,j_ + α_3_ * occasion_i,j_ | 5 | -737.25 | 1484.51 | 2.40 |
| α_0_ + α_1_ * season_i_ + α_2_ * effort_i,j_ + α_3_ * occasion_i,j_ + α_4_ * date_i,j_ | 6 | -736.72 | 1485.43 | 3.33 |
| **α_0_ + α_1_ * season_i_ + α_2_ * effort_i,j_** | **4** | **-738.77** | **1485.54** | **3.44** |
| α_0_ + α_1_ * season_i_ + α_2_ * effort_i,j_ + α_3_ * date_i,j_ + α_4_ * date^2^_i ,j_ | 6 | -737.17 | 1486.34 | 4.24 |

**Table 6.** Model selection table for the lowland tapir abundance sub-model where models are ranked by AIC from lowest to highest. Table includes state model specification, number of parameters (K), the log-likelihood (ℓ), and both AIC (2K - 2ℓ), and ΔAIC (AIC – lowest AIC). The top-ranked model after consideration of redundant parameters is highlighted in bold.

| **State model** | **Number of parameters (*K*)** | **Log-likelihood (ℓ)** | **AIC** | **ΔAIC** |
| --- | --- | --- | --- | --- |
| β_0_ + β_1_ * season_i_ + β_2_ * forest_i_ + β_3_ * latitude_i_ + β_4_ * river distance_i_ | 6 | -750.32 | 1512.65 | 0.00 |
| **β_0_ + β_1_ * season_i_ + β_2_ * forest_i_ + β_3_ * latitude_i_** | **5** | **-751.41** | **1512.83** | **0.18** |
| β_0_ + β_1_ * season_i_ + β_2_ * forest_i_ + β_3_ * latitude_i_ + β_4_ * longitude_i_ + β_5_ * river distance_i_ | 7 | -749.78 | 1513.55 | 0.90 |
| β_0_ + β_1_ * season_i_ + β_2_ * forest_i_ + β_3_ * latitude_i_ + β_4_ * longitude_i_ | 6 | -750.94 | 1513.88 | 1.24 |
| β_0_ + β_1_ * season_i_ + β_2_ * forest_i_ + β_3_ * latitude_i_ + β_4_ * elevation_i_ | 6 | -751.16 | 1514.32 | 1.67 |
| β_0_ + β_1_ * season_i_ + β_2_ * forest_i_ + β_3_ * latitude_i_ + β_4_ * river distance_i_ + β_5_ * soybean/corn_i_ | 7 | -750.27 | 1514.55 | 1.90 |
| β_0_ + β_1_ * season_i_ + β_2_ * forest_i_ + β_3_ * latitude_i_ + β_4_ * river distance_i_ + β_5_ * elevation_i_ | 7 | -750.31 | 1514.61 | 1.96 |
